# Molecular evolution of *DNMT1* in vertebrates: duplications in marsupials followed by positive selection

**DOI:** 10.1101/247643

**Authors:** David Alvarez-Ponce, María Torres-Sánchez, Felix Feyertag, Asmita Kulkarni, Taylen Nappi

**Affiliations:** Department of Biology, University of Nevada, Reno, Nevada, USA.; Department of Zoology and Physical Anthropology, Complutense University of Madrid, 28040 Madrid, Spain

**Keywords:** *DNMT1*, gene duplication, marsupials, wallaby, koala, Tasmanian devil, opossum

## Abstract

DNA methylation is mediated by a conserved family of DNA methyltransferases (Dnmts). The human genome encodes five Dnmts: Dnmt1, Dnmt2, Dnmt3a, Dnmt3b and Dnmt3L. Despite their high degree of conservation among different species, genes encoding Dnmts have been duplicated and/or lost in multiple lineages throughout evolution, indicating that the DNA methylation machinery has some potential to undergo evolutionary change. However, little is known about the extent to which this machinery, or the methylome, varies among vertebrates. Here, we study the molecular evolution of Dnmt1, the enzyme responsible for maintenance of DNA methylation patterns after replication, in 79 vertebrate species. Our analyses show that all studied species exhibit a single copy of *DNMT1*, with the exception of tilapia and marsupials (tammar wallaby, koala, Tasmanian devil and opossum), each of which exhibits two apparently functional *DNMT1* copies. Our phylogenetic analyses indicate that *DNMT1* duplicated before the divergence of marsupials (i.e., at least ~75 million years ago), thus giving rise to two *DNMT1* copies in marsupials (copy 1 and copy 2). In the opossum lineage, copy 2 was lost, and copy 1 recently duplicated again, generating three *DNMT1* copies: two putatively functional genes (copy 1a and 1b) and one pseudogene (copy 1ψ). Both marsupial copies (*DNMT1* copies 1 and 2) are under purifying selection, and copy 2 exhibits elevated rates of evolution and signatures of positive selection, suggesting a scenario of neofunctionalization. This gene duplication might have resulted in modifications in marsupial methylomes and their dynamics.

## INTRODUCTION

In vertebrate genomes, cytosine methylation is widespread (e.g., 60–90% of CpGs are methylated in mammals [1, 2]) and plays pivotal roles in the silencing of gene expression and transposable elements, gene imprinting, and X-chromosome inactivation [3]. DNA methylation is mediated by a conserved family of DNA methyltransferases (Dnmts). The human genome encodes five members of this family: Dnmt1, Dnmt2, Dnmt3a, Dnmt3b and Dnmt3L. Dnmt3a and Dnmt3b are responsible for *de novo* DNA methylation in germ cells and early embryos [4, 5]. An additional member of the Dnmt3 group, Dnmt3L, does not exhibit catalytic activity, but acts as a regulator of Dnmt3a and Dnmt3b. Once established by Dnmt3a and Dnmt3b, methylation patterns are maintained by Dnmt1, which copies them to the daughter DNA strand after replication [6]. Despite their sequence and structural similarity to Dnmt1 and Dnmt3s, Dnmt2 methylates the anticodon loop of aspartic acid transfer RNA, rather than DNA [7, 8].

Prior comparative analyses of distantly related organisms have revealed a number of gene duplications and losses in the evolutionary history of the genes encoding Dnmts. A number of organisms lack such genes (and DNA methylation), including the yeast *Saccharomyces cerevisiae* and the nematode *Caenorhabditis elegans*, and the number of Dnmts of each kind varies among lineages [2, 9 10 11 12 13 14]. For instance, *DNMT3C*, a mouse retrogene that evolved by duplication of *DNMT3B*, has been recently shown to be responsible for silencing young retrotransposons in the male germ line [15]. All three *DNMT* classes present in animals (classes 1, 2 and 3) are duplicated in some insect groups and completely absent from others [13]. Some insects, including Diptera, have lost DNA methylation, and insects with a methylome include some lacking Dnmt1s or Dnmt3s, indicating that neiher of the enzymes individually is essential for DNA methylation [13].

Little is known about the extent to which the DNA methylation machinery, or the methylome, may vary among vertebrates, and particularly among mammals. Molecular evolution studies of the DNA methylation machinery in vertebrates include some comparative analyses of members of the Dnmt3 group [16, 17], but less is known about the evolution of *DNMT1*. The human *DNMT1* gene has 40 exons and encodes a full, 1616-amino acid somatic isoform (Dnmt1s) and a truncated isoform expressed in oocytes (Dnmt1o), which lacks the first 118 amino acids. Dnmt1 proteins contain an N-terminal regulatory region and a C-terminal catalytic domain, separated by a KG repeat. In the Dnmt1s isoform, the regulatory region comprises a DNA methyltransferase associated protein (DMAP) binding domain, a nuclear localization signal (NLS), a replication foci targeting sequence (RFTS), a cysteine-rich DNA binding domain (CXXC), an autoinhibitory linker that prevents *de novo* methylation, and two bromo-adjacent homology domains (BAH1 and BAH2), among other protein-interaction domains (Fig. 1; for a comprehensive review, see ref. 18). A direct interaction between the N-terminal and the C-terminal domains seems to be necessary for enzyme activation [19]. Activated Dnmt1 shows high affinity for hemimethylated CG sites. *DNMT1*-null mouse embryos die soon after implantation and exhibit delayed development and structural abnormalities [20], and overexpression of *DNMT1* has been observed in multiple cancer tissues [21 22 23 24].

**Fig. 1.**
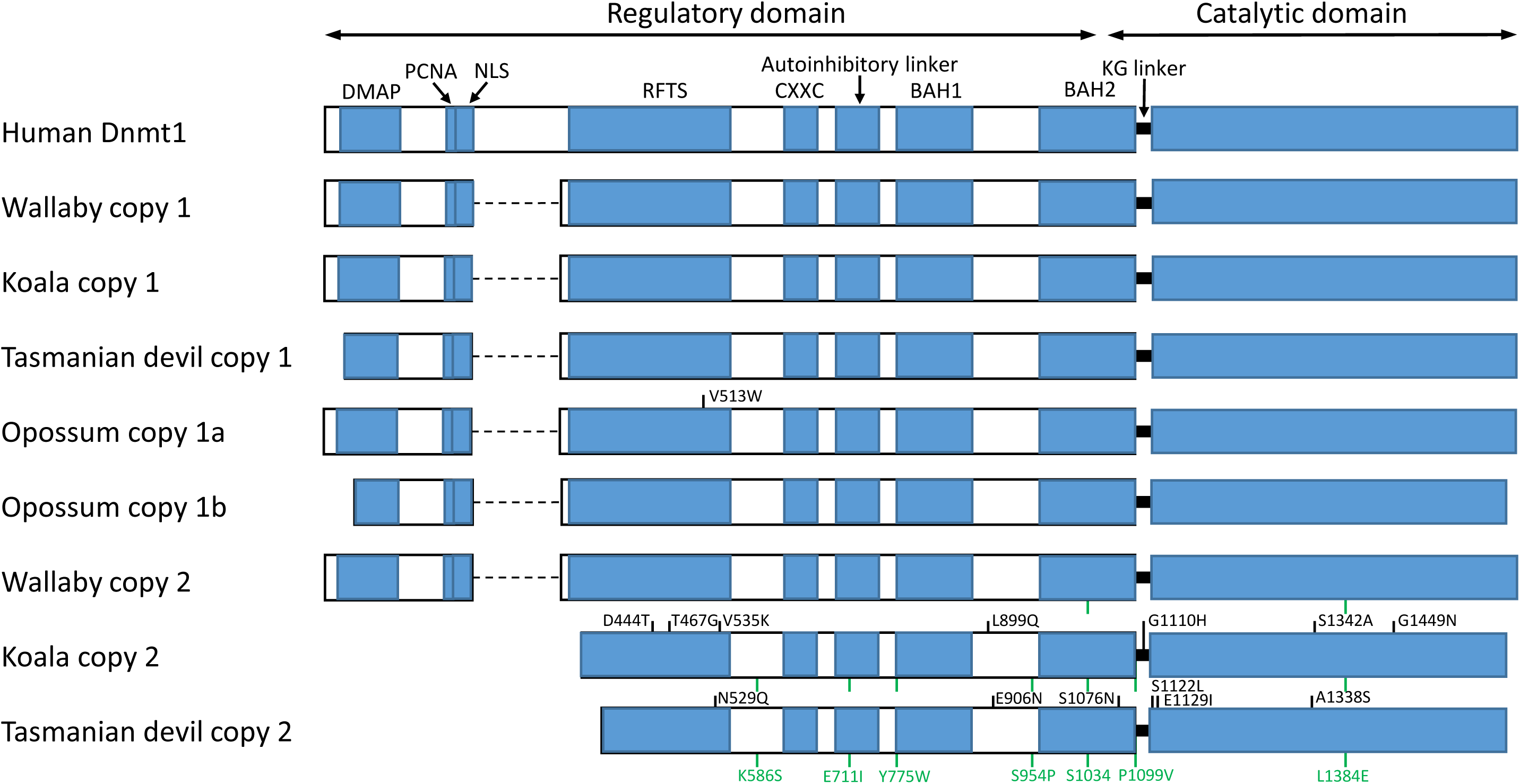
Structure of Dnmt1 proteins in human and in marsupials. The human Dnmt1s isoform is represented. Sites under positive selection specific to one of the sequences are represented in black. Sites under positive selection shared across multiple sequences (due to positive selection in an internal branch) are represented in green, and their coordinates are only indicated for the last sequence. Amino acid coordinates refer to the human protein. Dashed lines represent missing parts. DMAP, DNA methyltransferase associated protein-binding domain; PCNA, proliferating cell nuclear antigen-binding domain; NLS, nuclear localization signal; RFTS, replication foci targeting sequence; CXXC, cysteine-rich DNA binding domain; BAH, bromo-adjacent homology domains.

Here, with the aim of identifying potential differences among the methylation machineries of verterates, we study the molecular evolution of *DNMT1* in 79 vertebrate species. Our analyses reveal that all studied species exhibit a single *DNMT1*, with the only exception of tilapia and marsupials (tammar wallaby, koala, Tasmanian devil and opossum), each of which exhibit two putatively functional *DNMT1* copies. Our phylogenetic analyses indicate that *DNMT1* duplicated before the divergence of marsupials (at least ~75 million years ago), thus giving rise to two *DNMT1* copies (copies 1 and 2) in marsupials. Copy 2 was subsequently lost in the opossum lineage, whereas copy 1 recently duplicated again twice in the opossum lineage, to generate three genes in this species: two putatively functional ones (copies 1a and 1b) and one pseudogene (copy 1ψ). Both marsupial copies (*DNMT1* copies 1 and 2) are under purifying selection, and copy 2 exhibits signatures of positive selection, suggesting a scenario of neofunctionalization. We discuss how the presence of two DNMT1s in marsupials might have affected their methylome.

## RESULTS

### *DNMT1* duplicated in a marsupial ancestor and one of the resulting copies further duplicated in an opossum ancestor

We searched the complete genomes of 58 mammals, 5 birds, 2 reptiles, one amphibian and 13 fish (Table S1) for orthologs of the human *DNMT1* gene. The studied mammalian species included 53 placentarians, four marsupials (tammar wallaby [25], koala [26], Tasmanian devil [27] and opossum [28]) and one monotreme (platypus [29]) (Table S1). All studied genomes exhibit a single *DNMT1* copy, with the exception of tilapia and the four marsupials, each of which exhibits two putatively functional copies. In addition, 8 of the studied genomes (including opossum) exhibit pseudogenes maintaining homology to a substantial length of *DNMT1*.

According to the annotations of the Ensembl database [30], the tilapia genome contains two DNMT1 copies (ensemble gene IDs: ENSONIG00000001574 and ENSONIG00000007221). The first copy encodes a full Dnmt1 protein (1505 amino acids). The second copy is located in a very small scaffold (AERX01074151.1, 3084 nucletides), which only covers exons 36–40 (184 amino acids). These exons are identical between both copies, but many differences (single-point mutations and indels) are observed in the introns. These observations indicate a very recent duplication of *DNMT1* in tilapia, but the fact that only a small portion of one of the copies is available prevents further analysis. Thus, it cannot be discarded that one of the copies is a pseudogene, or in the process of pseudogenization.

Some of the *DNMT1* copies identified were unannotated, or their exon/intron structure was incorrectly annotated in the Ensembl [30] and nr databases. Where necessary, marsupial and platypus sequences were re-annotated manually using the human *DNMT1* as reference (see Methods), and incomplete sequences (due to their location in partially sequenced genomic regions) were completed using available RNA-seq data [31 32 33]. The resulting protein sequences are shown in Figs. 2 and 3.

**Fig. 2.**
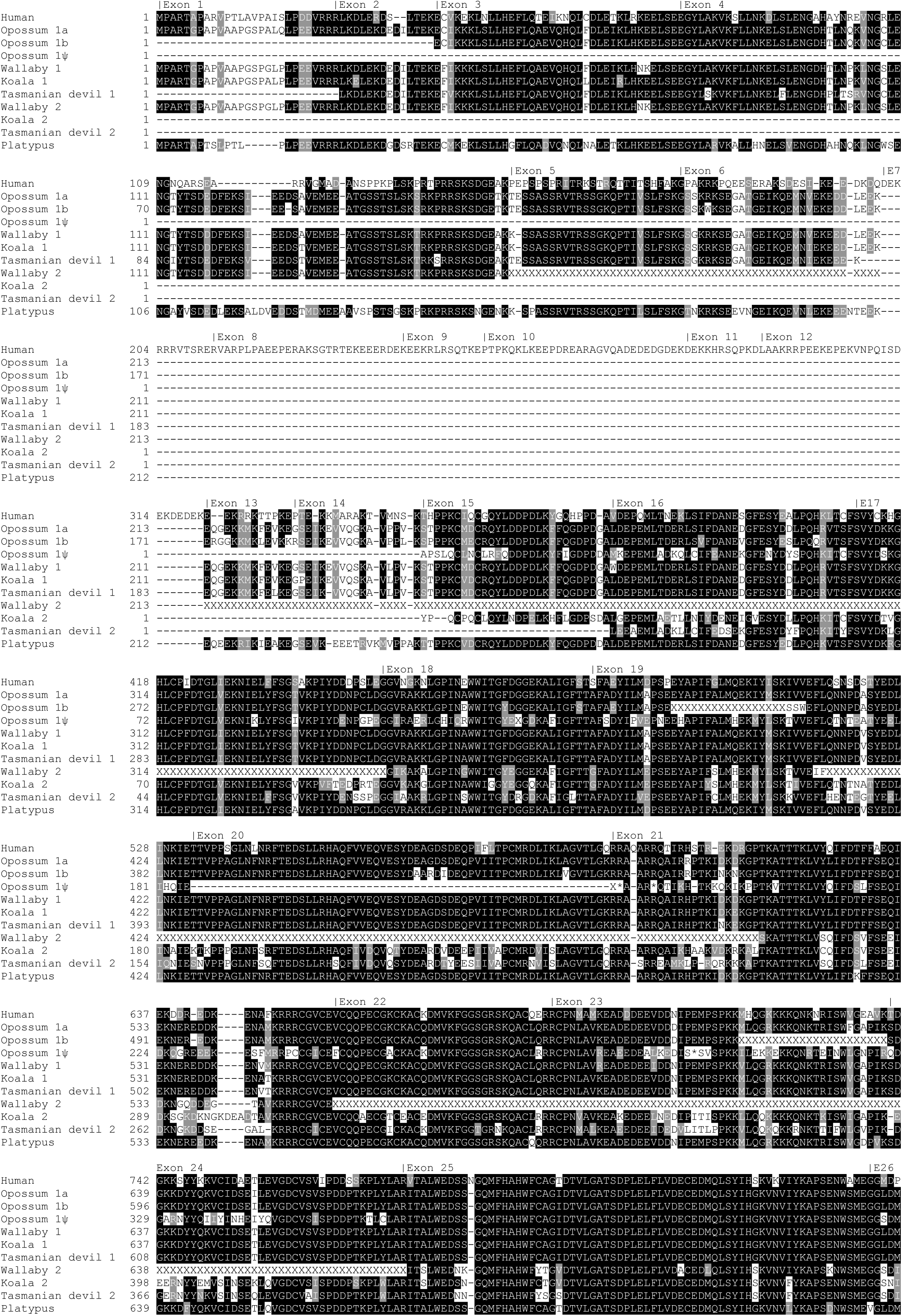
Alignment for the N-terminal part of Dnmt1 in human marsupials and platypus. The human sequence corresponds to the Dnmt1s isoform. Dashes represent alignment gaps or missing regions. Stretches of “X” symbols represent unsequenced regions. Single “X” symbols represent incomplete codons (e.g., due to frameshift mutations).

**Fig. 3.**
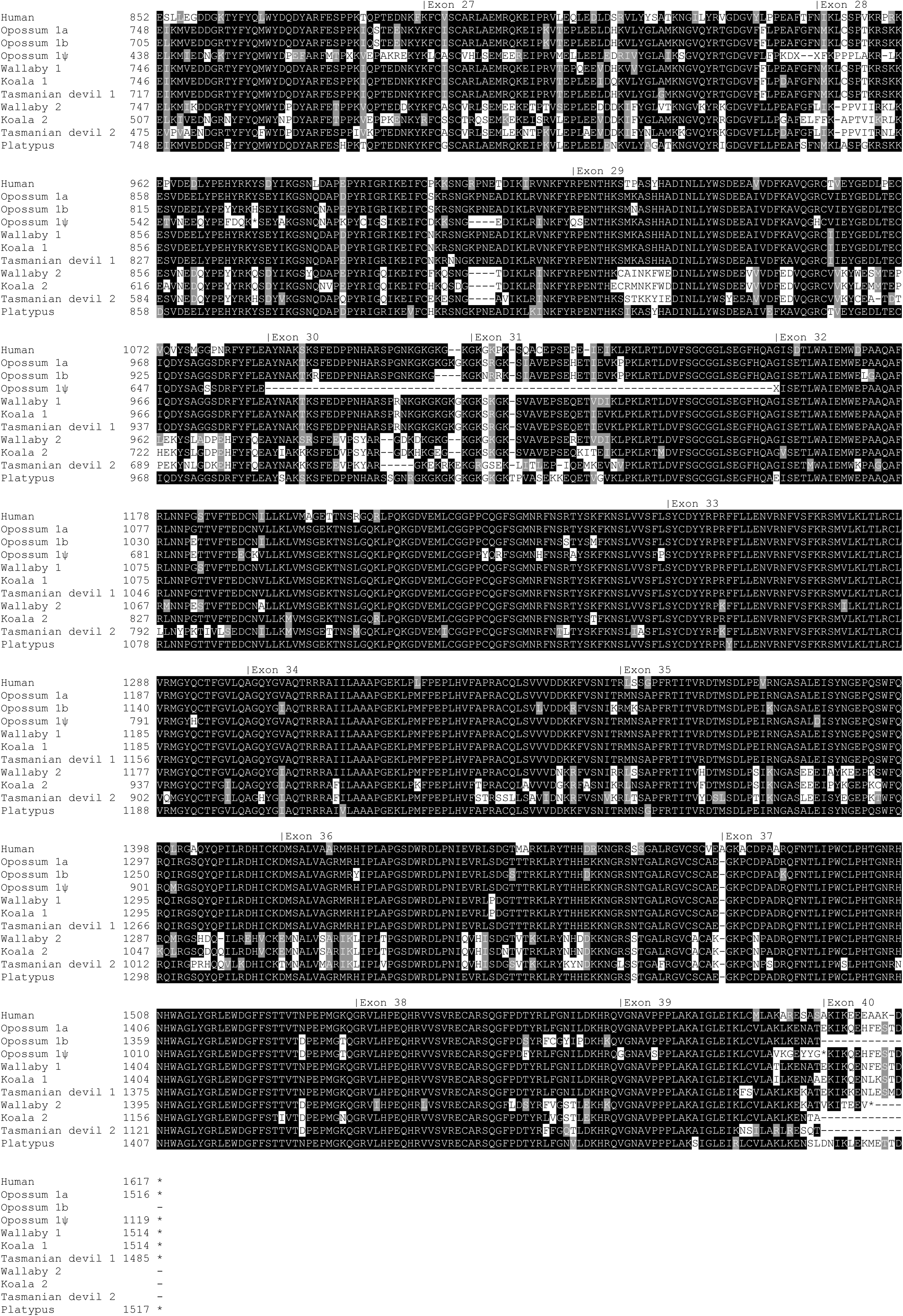
Alignment for the C-terminal part of Dnmt1 in human, marsupials and platypus. The human sequence corresponds to the Dnmt1s isoform. Dashes represent alignment gaps or missing regions. Stretches of “X” symbols represent unsequenced regions. Single “X” symbols represent incomplete codons (e.g., due to frameshift mutations).

In the case of opossum, the three *DNMT1* copies (two putatively functional genes and one pseudogene) are located in tandem in chromosome 3, suggesting two recent duplication events. The two koala sequences are also located in the same scaffold (NW_018344010.1, 26.8 Kb apart). Tasmanian devil’s scaffold GL841404.1 contains one of the copies and part (exons 37–39) of the other copy, 4.6 Kb apart; the other exons of the second copy are located in another two contigs, most likely due to assembly errors (see Methods). The wallaby copies are located in different scaffolds (GeneScaffold_10206 and GeneScaffold_8347); however, these scaffolds are small (45.9 and 90.7 Kb, respectively), and therefore we cannot discard the possibility that both wallaby copies are also closely linked (Table S1).

The three opossum copies exhibit high sequence similarity (copy 1a vs. copy 1b: *d*_N_ = 0.022; *d*_S_ = 0.047; copy 1a vs. copy 1ψ: *d*_N_ = 0.081; *d*_S_ = 0.201; copy 1b vs. copy 1ψ: *d*_N_ = 0.091; *d*_S_ = 0.205; measures of divergence calculated using the Nei-Gojobori method [34] and the Jukes-Cantor correction [35] as implemented in DnaSP version 5.10.01 [36]), whereas the wallaby, koala and Tasmanian devil copies are much more divergent (wallaby’s copy 1 vs. copy 2: *d*_N_ = 0.114; *d*_S_ = 0.440; koala’s copy 1 vs. copy 2: *d*_N_ = 0.120; *d*_S_ = 0.395; Tasmanian devil’s copy 1 vs. copy 2: *d*_N_ = 0.146; *d*_S_ = 0.521) (Figs. 2 and 3). These observations, combined with the results of our phylogenetic analysis (Fig. 4), and the known marsupial phylogeny (among the studied species, wallaby and koala are the most closely related, followed by Tasmanian devil and opossum [37, 38]), suggest a scenario in which: (a) *DNMT1* duplicated in a common ancestor of marsupials, giving rise to copies 1 and 2; (b) copy 2 was lost from the opossum lineage; and (c) copy 1 was recently duplicated twice in the opossum lineage, giving rise to two putatively functional copies and one pseudogene. The relative order of the latter two events is unclear. Based on this inferred scenario, we named the three opossum copies as copy 1a (chromosome 3, positions 431,108,118–431,161,113), copy 1b (positions 431,298,625–431,342,040) and copy 1ψ (pseudogene, positions 431,228,446–431,291,545). Copy 1a was already reported by Ding et al. [39], and the presence of a second copy in opossum was noted by Nikkelsen et al. [28].

**Fig. 4.**
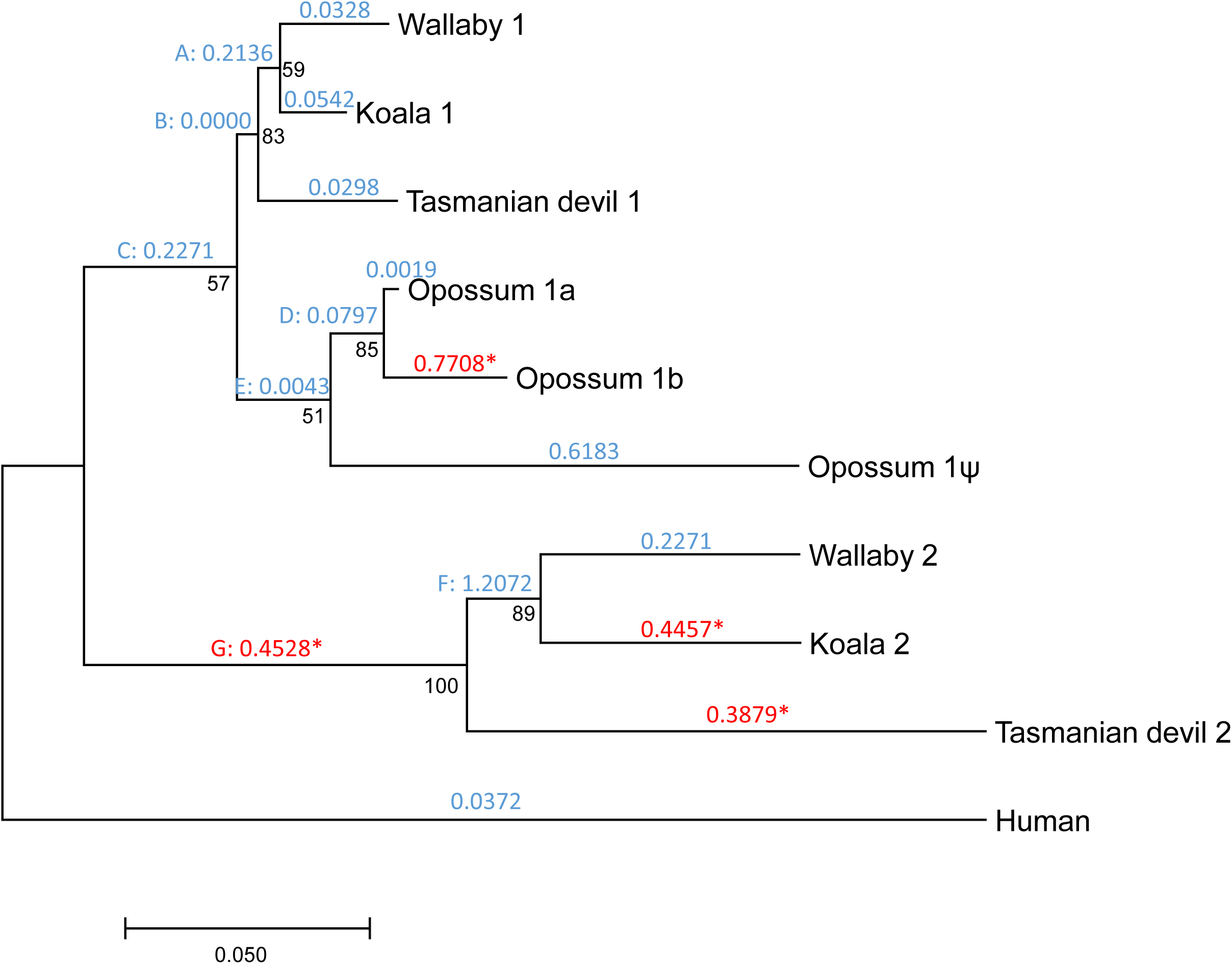
Phylogenetic tree showing the duplication of *DNMT1* in marsupials. Numbers in black represent bootstrap values. Numbers in blue or red above each branch represent *d*_N_/*d*_S_ values according to the free-ratios model. For branches under positive selection according to the branch-site test, *d*_N_/*d*_S_ ratios are represented in red and are followed by an asterisk. Internal branches are labelled with capitals letters.

All marsupial and monotreme sequences lack exons 7–12 (using the human Dnmt1s isoform as reference), consistent with the opossum sequence reported by Ding et al. [39] (Fig. 1). A BLASTP search (*E-value* < 10^−3^) against all proteomes available in the Ensembl database failed to find any significant hit in non-placentarians, indicating that these exons, which encode amino acids 201–320, were acquired in placentarians. These amino acids overlap with the following regions: region of interaction with the PRC2/EED-EZH2 complex (amino acids 1–606), region of interaction with Dnmt3b (positions 149–217), NLS (positions 177–205) and homodimerization region (positions 310–502). In addition, koala’s copy 2 lacks exons 1–14 (first 347 amino acids), and Tasmanian devil’s copy 2 lacks exons 1–15 (amino acids 1–374). Thus, the encoded proteins lack the regions of interaction with DMAP (positions 18–103), Dnmt3a (positions 1–148), Dnmt3b (positions 149–217), and PCNA (positions 163–174), the NLS (positions 177–205), part of the homodimerization (positions 310–502) and RFTS (positions 331–550) regions, and the region of interaction with the PRC2/EED-EZH2 complex (positions 1–606). Nonetheless, all marsupial and monotreme Dnmt1s appear to include a complete CXXC domain, an autoinhibitory linker, the BAH1 and BAH2 domains, and the catalytic domain (Fig. 1), thus being potentially functional. The opossum pseudogene (copy 1ψ) lacks exons 1–16, 20, and 30–31, and contains five stop codons (two in exon 21, one in exon 23, one in exon 28, and one in the codon shared between exons 39 and 40) and two frameshift mutations (exons 18 and 26).

### Marsupial *DNMT1* copies are differentially expressed

We next attempted to determine in which tissues, and to what extent, each copy is expressed. First, we searched the transcriptomes of a number of koala tissues [40] for transcripts corresponding to copy 1 and copy 2, finding only transcripts for copy 1. Second, we searched two Tasmanian devil transcriptomic datasets (lymph and spleen) for sequences similar to *DNMT1*, finding only reads for copy 1. Third, we mined RNA-seq data for 5 wallaby tissues (testes, male liver, female liver, male blood and female blood; ref. 32), and identified 11,267 reads specific to copy 1 and only 5 reads specific to copy 2 (another 735 reads matched both copies; Table 1). Finally, we mined RNA-seq data for 11 opossum tissues (testis and male and female brain, cerebral cortex, heart, kidney and liver; ref. 31). A total of 3831, 194 and 290 reads matched opossum’s copies 1a, 1b and 1ψ, respectively (Table 2).

**Table 1.** Number of RNA-seq reads matching wallaby’s copies 1 and 2.

| **Tissue** | **Run accession number** | **Copy 1** | **Copy 2** |
| --- | --- | --- | --- |
| Male liver | SRR1041778 | 502 | 0 |
| Female liver | SRR1552212 | 1340 | 1 |
| Male blood | SRR1552202 | 371 | 0 |
| Female blood | SRR1552210 | 233 | 2 |
| Testis | SRR1041779 | 8821 | 2 |
| Total: |  | 11267 | 5 |

**Table 2.** Number of RNA-seq reads matching opossum’s copies 1a, 1b and 1ψ.

| **Tissue** | **Run accession number** | **Copy 1a** | **Copy 1b** | **Copy 1ψ** |
| --- | --- | --- | --- | --- |
| Male brain | SRR306744 | 158 | 4 | 11 |
| Male cerebral cortex | SRR306746 | 279 | 9 | 37 |
| Male heart | SRR306750 | 318 | 8 | 14 |
| Male kidney | SRR306752 | 190 | 40 | 40 |
| Male liver | SRR306754 | 87 | 1 | 8 |
| Testis | SRR306756 | 1739 | 52 | 24 |
| Female brain | SRR306743 | 106 | 5 | 11 |
| Female cerebral cortex | SRR306745 | 265 | 20 | 51 |
| Female heart | SRR306748 | 142 | 5 | 3 |
| Female kidney | SRR306751 | 344 | 45 | 83 |
| Female liver | SRR306753 | 203 | 5 | 8 |
| Total: |  | 3831 | 194 | 290 |

### Both marsupial *DNMT1* copies are under purifying selection

We used PAML [41] to estimate the non-synonymous to synonymous divergence ratio (*d*_N_/*d*_S_) in each of the branches of the gene tree. We restricted this analysis to human and the four marsupials, as incomplete genomic data and annotation errors in many of the other species would have hindered our analyses. This ratio was substantially below one in all branches of the phylogeny, except in the internal branch leading to the most recent common ancestor (MRCA) of wallaby’s and koala’s copy 2 (Fig. 4). This indicates that nonsynonymous changes are under substantial purifying selection in all the sequences studied, suggesting that all copies are functional, or that they pseudogenized only recently – which is the case for opossum’s copy 1ψ (*d*_N_/*d*_S_ = 0.618).

The *d*_N_/*d*_S_ ratios varied substantially among the different branches (Fig. 4). Indeed, the free-ratios model fit the data significantly better than the one-ratio model M0 (2Δ*ℓ* = 438.07, *P* = 3.71×10^−83^), indicating significant heterogeneity in the *d*_N_/*d*_S_ ratios. Remarkably, *d*_N_/*d*_S_ was substantially higher in copy 2 than in copy 1 (Fig. 4). In addition, *d*_N_/*d*_S_ was 0.0019 in the branch leading to opossum’s copy 1a, and 0.7708 in the branch leading to opossum’s copy 1b. This increase in the *d*_N_/*d*_S_ ratios of copy 2 (wallaby, koala and Tasmanian devil) and copy 1b (opossum) could be explained by a relaxation of purifying selection acting on protein sequences and/or by positive selection in these copies.

### Marsupials’ copy 2 of *DNMT1* is under positive selection

We then used PAML to test for signatures of positive selection. The M8 vs. M7 test was significant (2Δ*ℓ* = 8.36, *P* = 0.015), indicating that a fraction of codons were under positive selection. We then used a branch-site test (model A vs. null model A1; refs. 42, 43) to infer the action of positive selection at each of the branches of the phylogeny, except the branch leading to the opossum pseudogene. The test was significant for the external branches leading to koala’s copy 2, Tasmanian devil’s copy 2, and opossum’s copy 1b, and for the internal branch leading to the MRCA of the copy 2 of wallaby, koala and Tasmanian devil. The *d*_N_/*d*_S_ values for these branches are represented in red and marked with an asterisk in Fig. 4, and more detailed results are provided in Table 3.

**Table 3.** Branch-site tests of positive selection.

| Branch<sup>a</sup> | Log-likelihood<br>model A | Log-likelihood<br>null model A1 | $2\Delta\ell$ | *P*-value | $\omega_s$<sup>b</sup> | Selected codons<sup>b</sup> |
| --- | --- | --- | --- | --- | --- | --- |
| Wallaby 1 | -10,537.29 | -10,537.29 | $5.2 \times 10^{-4}$ | 0.497 | 1.000 | |
| Koala 1 | -10,537.29 | -10,537.29 | 0.00 | 0.500 | 1.000 |  |
| Tasmanian devil 1 | -10,537.29 | -10,537.29 | 0.00 | 0.500 | 1.000 |  |
| Opossum 1a | -10,537.29 | -10,537.29 | 0.00 | 0.500 | 1.000 |  |
| Opossum 1b | -10,527.28 | -10,531.13 | 7.70 | 0.003* | 11.328 | V513W |
| Wallaby 2 | -10,535.71 | -10,535.71 | 0.00 | 0.500 | 1.000 |  |
| Koala 2 | -10,526.31 | -10,527.99 | 3.35 | 0.034* | 2.960 | T467G, S1342A, G1449N |
| Tasmanian devil 2 | -10,517.81 | -10,523.87 | 12.12 | $2.5 \times 10^{-4}$*** | 5.101 | E906N, S1076N, A1338S |
| Human | -10,537.12 | -10,537.29 | 0.34 | 0.279 | 43.846 |  |
| A | -10,537.29 | -10,537.29 | 0.00 | 0.500 | 1.000 |  |
| B | -10,537.29 | -10,537.29 | 0.00 | 0.500 | 1.000 |  |
| C | -10,537.13 | -10,537.13 | 0.00 | 0.500 | 1.000 |  |
| D | -10,537.29 | -10,537.29 | 0.00 | 0.500 | 1.000 |  |
| E | -10,537.13 | -10,537.13 | 0.00 | 0.500 | 1.000 |  |
| F | -10,532.75 | -10,533.68 | 1.85 | 0.087 | 150.054 |  |
| G | -10,526.76 | -10,529.87 | 6.22 | 0.006* | 3.717 | S1034, L1384E |

<sup>a</sup>Internal branches are represented with letters as in Fig. 4.
<sup>b</sup> $d_N/d_S$ for the class of codons under positive selection.
<sup>c</sup>For each mutation, the first letter and the number correspond to the amino acid in human DNMT1s, and the last letter corresponds to the mutation observed in the sequence(s) of interest. For the mutation S1034, the final amino acid is not provided because it is not the same in all the descendants of branch G.
\*, $P < 0.05$ ; \*\*\*, $P < 0.001$ .

A total of 9 codons were detected to be under positive selection: one in the opossum’s copy 1b, three in koala’s copy 2, three in Tasmanian devil’s copy 2, and 2 in the internal branch leading to the MRCA of copy 2 of wallaby, koala and Tasmanian devil. Sites under positive selection were different in each branch, and affected the catalytic domain (4 codons), the site of interaction with the PRC2/EED-EZH2 complex (6 codons), the BAH2 domain (2 codons) and the homodimerization domain (1 codon; Fig. 1; Table 3).

### Reanalysis removing incomplete sequences

The marsupial *DNMT1* coding sequences (CDSs) used in this study are complete or almost complete (Figs. 2 and 3). The only notable exceptions are wallaby’s copy 2, for which 409 codons remain unsequenced due to limited genome coverage (2×; ref. 25), and the opossum pseudogene, which lacks 19 exons. This means that our natural selection analyses were limited to only 826 codons. We repeated our analysis after removing these sequences from our analysis, rendering 1172 codons analyzable(present in all sequences). We obtained similar results: First, the *d*_N_/*d*_S_ ratio was substantially higher in copy 2 than in copy 1, and in opossum’s copy 1b (*d*_N_/*d*_S_ = 0.808) than in opossum’s copy 1a (*d*_N_/*d*_S_ = 0.000; Fig. 5). Second, positive selection was detected in the external branches leading to opossum’s copy 1b, koala’s copy 2 and Tasmanian devil’s copy 2, and in the internal branch leading to the MRCA of koala’s copy 2 and Tasmanian devil’s copy 2. This analysis detected a total of 21 codons under positive selection (including the 9 ones detected before), which affected the catalytic domain (4 codons), the site of interaction with the PRC2/EED-EZH2 complex (10 codons), the BAH2 domain (3 codons), the homodimerization domain (2 codons), the autoinhibiroty linker (1 codon), and the KG linker (1 codon; Fig. 1; Table S1).

**Fig. 5.**
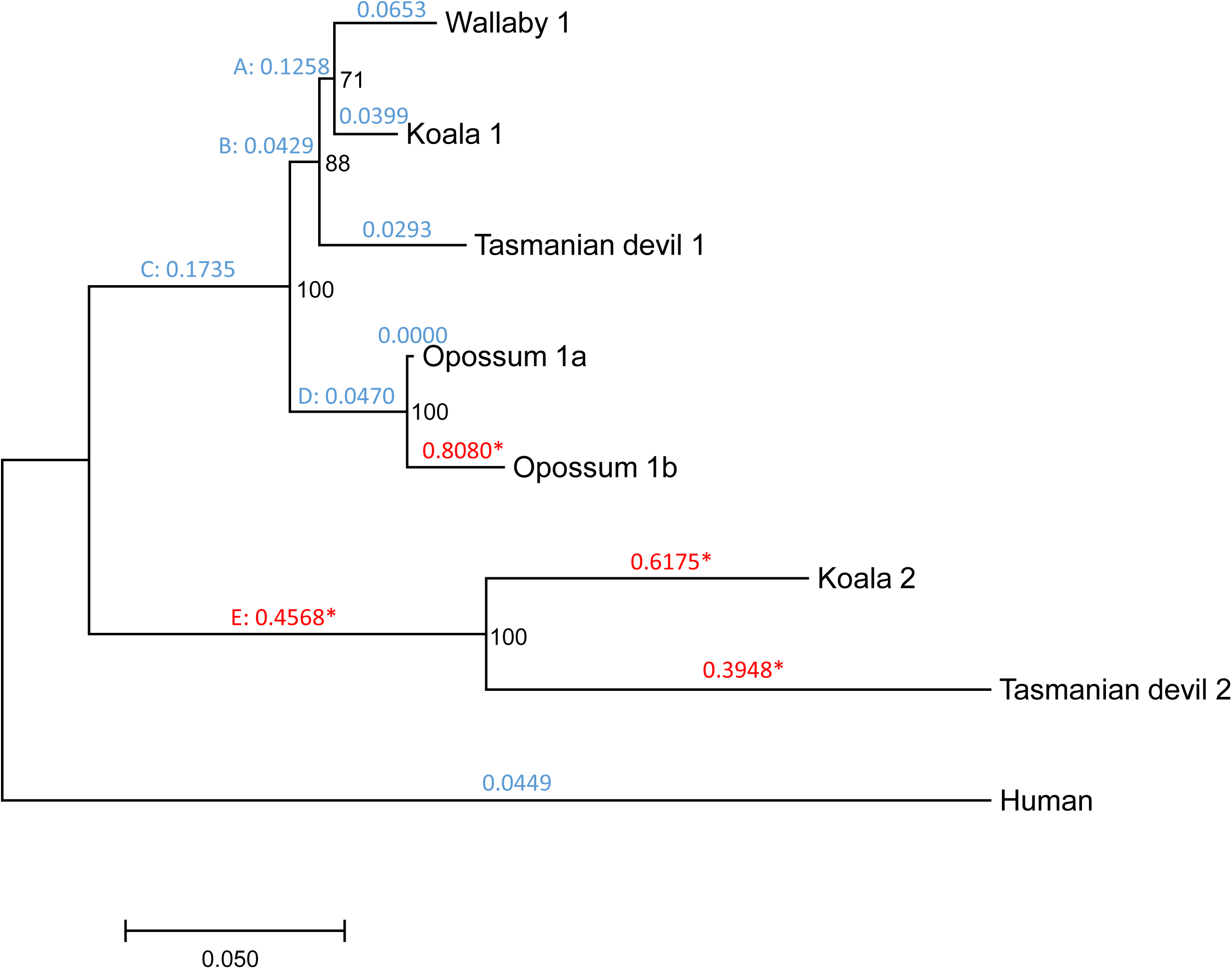
Phylogenetic tree showing the duplication of *DNMT1* in marsupials, removing wallaby’s copy 2 and opossum’s copy 1ψ. Numbers in black represent bootstrap values. Numbers in blue or red above each branch represent *d*_N_/*d*_S_ values according to the free-ratios model. For branches under positive selection according to the branch-site test, *d*_N_/*d*_S_ ratios are represented in red and are followed by an asterisk. Internal branches are labelled with capitals letters.

## DISCUSSION

Our analyses indicate that the *DNMT1* gene duplicated in a common ancestor of marsupials, giving rise to two copies (copies 1 and 2). The opossum lineage and the wallaby/koala/Tasmanian devil lineage diverged ~75 million years ago [37, 38], implying that the *DNMT1* duplication occurred prior to that time. Copy 2 was subsequently lost in the opossum lineage. Copy 2 is expressed at very low, or even undetectable levels, at least in the wide range of wallaby (Table 1), koala [40] and Tasmanian devil tissues examined. However, both copies exhibit *d*_N_/*d*_S_ ratios lower than one, and none exhibit signatures of pseudogenization (premature stop codons or frameshift mutations) indicating that they are expressed—perhaps in tissues not included in our analyses, in early developmental stages or under certain environmental conditions— and functional. Otherwise, signatures of pseudogenization and a *d*_N_/*d*_S_ close to 1 would be expected. Part of the regulatory region of koala’s and Tasmanian devil’s copy 2 appear to have been lost; however, all *DNMT1* copies retain the catalytic domain and a significant fraction of the regulatory region, suggesting that they are functional—of note, the human DNMT1o isoform is functional despite also lacking part of the regulatory region.

Remarkably, copy 2 exhibits a high *d*_N_/*d*_S_ ratio compared to copy 1, in addition to signatures of positive selection. These results suggest a scenario of neofunctionalization, in which copy 1 may have retained the function of the ancestral *DNMT1*, and copy 2 may have acquired a new or modified function. Signatures of positive selection can be detected in the branch leading to the MRCA of wallaby’s, koala’s, and Tasmanian devil’s copy 2, and in the external branches leading to koala’s and Tasmanian devil’s copy 2 (Figs. 4 and 5; Tables 3 and 4). These observations indicate that neofunctionalization occurred both before and after the divergence of wallaby, koala and Tasmanian devil (i.e., both before and after ~60 million years ago; refs. 37, 38). Substitutions under positive selection affect different domains, making it difficult to predict how they may have affected the function of copy 2.

Copy 1 recently underwent another two duplication events in the opossum lineage, which resulted in three genes (copies 1a, 1b and the pseudogene 1ψ) located in tandem in chromosome 3. Their high degree of similarity, along with our phylogenetic analyses (Fig. 4), indicate that these sequences are the result of the duplication of the copy 1 of *DNMT1*, and that they are not remnants of the ancestral duplication identified in the other marsupials. Opossum’s copy 1b also exhibits an elevated *d*_N_/*d*_S_ (compared to copy 1b) and signatures of positive selection, which would also suggest neofunctionalization in the copy 1b. However, in this case we are skeptical about our inference of positive selection, because the only codon inferred to be under positive selection with high probability (V513 in the human protein, a tryptophan in opossum’s copy 1b) is located near an unsequenced region of the opossum genome (Fig. 2), and such regions are prone to sequencing errors. Opossum’s copy 1b is expressed at lower levels than copy 1a in the tissues included in our analyses (Table 2).

It is currently not possible to infer the functions of marsupial *DNMT1* derived duplicates (copy 2 of wallaby, koala and Tasmanian devil and copy 1b of opossum). We propose three different possible scenarios. First, as both marsupial *DNMT1* copies seem to be expressed in different sets of tissues (Tables 1 and 2), positive selection in the derived *DNMT1* copies may simply reflect subtle adjustments to the biochemistry of the tissue or tissues in which they are expressed. Second, assuming that the function of both marsupial *DNMT1* copies is similar to that of the ancestral *DNMT1*—maintenance of methylation patterns throughout the life of the animal after each DNA replication event— it is possible that an increased Dnmt1 abundance may cause marsupial methylomes to be particularly stable during aging—in other mammals methylation patterns change during the lifespan of an organism [44]. This, however would only apply to the unknown tissue or tissues (or developmental stages or environmental conditions) in which the derived copies are expressed at substantial levels. Third, the duplication of *DNMT1* may have caused marsupial genomes to be hypermethylated. Given that methylated cytosines have an increased mutation rate [45], this scenario might explain the low GC content of marsupial genomes [25, 27, 28, 46]. However, this scenario would require that the derived *DNMT1* copies would act as *de novo* DNMTs rather than maintenance DNMTs, which is at odds with the presence of an autoinhibitory linker in the proteins encoded by both copies. Additional functional studies of marsupial Dnmt1s, and methylome data for marsupials —which is currently unavailable— will be required to establish their functions.

## CONCLUSIONS

Our analyses of 79 vertebrate genomes reveal that all studied species exhibit a single DNMT1, with the exception of tilapia and marsupials (wallaby, koala, Tasmanian devil and opossum), each of which exhibit two apparently functional *DNMT1* copies. Our phylogenetic analyses indicate that *DNMT1* duplicated before the divergence of marsupials (at least ~75 million years ago), thus giving rise to *DNMT1* copies 1 and 2. Copy 2 was lost in the opossum lineage, and copy 1 recently duplicated again to generate three opossum genes: two putatively functional ones and one pseudogene. Both *DNMT1* copies are under purifying selection, and copy 2 is under positive selection. These results suggest a scenario of neofunctionalization.

## METHODS

### Gene identification and annotation

In order to identify *DNMT1* orthologs in the studied vertebrate genomes, we conducted TBLASTN searches against the Ensembl database (release 90; ref. 30), using the human Dnmt1s protein sequence as query and an *E-value* cut-off of 10^−10^. The koala genome was queried in the nr database, as it is not represented in Ensembl. Only scaffolds with at least 450 identities (added across the different TBLASTN hits) were considered.

Where necessary, wallaby, koala, Tasmanian devil, opossum and platypus sequences were manually re-annotated using the intron/exon structure of human *DNMT1* as reference. For that purpose, incorrectly annotated exons (those not showing significant similarity to the human sequence) were removed, and missing exons were searched for using TBLASTN and BLASTN searches. Putative stop codons and frameshift mutations were confirmed by visualization of the corresponding original reads in the trace archive database.

In the case of Tasmanian devil’s copy 2 and platypus’ *DNMT1*, exons present in different scaffolds were combined into a single gene annotation. The platypus *DNMT1* exons are distributed along two small contigs: Contig12710 (18.1 Kb) and Contig19880 (17.7 Kb) (Table S1). In the current Tasmanian devil assembly, the exons of copy 2 are distributed across three different scaffolds: exons 16–24 are located in scaffold GL841374.1 (4.0 Mb), exons 25–36 are located in GL843446.1 (17.2 Kb), and exons 37–39 are locate in GL841404.1 (1.6 Mb); this is probably the result of assembly errors.

Some of the exons of wallaby’s copies 1 and 2, opossum’s copy 1b, and the single copy of platypus, could not be recovered (or completely recovered) from available genome assemblies because they were located in unsequenced regions. We thus attempted to recover these exons from available RNA-seq datasets [31-33]. In the case of wallaby’s copy 2, this was not possible due to the very few reads available (Table 1), and in the case of opossum’s copy 1b it was not possible either due to the high similarity between copies 1a and 1b.

### Gene expression levels in different tissues

We used koala’s copy 2 as query in a TBLASTN search against the koala transcriptome [40]; all retrieved copies, however, corresponded to copy 1. Similarly, we used Tasmanian devil’s copy 2 as query in a TBLASTN search against all the RNA-seq reads available for two Tasmanian devil tissues (lymph and spleen; SRA accession numbers: ERR695583 and ERR695584), finding again only reads corresponding to copy 1.

We next mined RNA-seq datasets for a number of tissues of wallaby [32] and opossum [31], in order to measure expression levels of each of the *DNMT1* copies in the different tissues. For each read, it was determined whether it perfectly matched (it was contained in) one or more of the copies in the genome of interest, using an in-house PERL script. Reads that matched more than one copy were not used to compute expression levels.

### Phylogenetic analyses

The CDSs of human, wallaby, koala, Tasmanian devil, opossum and platypus were translated in silico into protein sequences. The protein sequences were aligned using ProbCons version 1.12 [47], and the resulting sequences were used to guide the alignment of the CDSs. Alignments were visualized and, where necessary, manually edited using BioEdit version 7.2.5 [48]. A phylogenetic tree was obtained using the maximum-likelihood method implemented in MEGA7 [49], using the Tamura-Nei model [50] and 1000 bootstraps.

### Natural selection analyses

The codeml program in the PAML package, version 4.4d [41] was used to conduct natural selection analyses. The free-ratios model was used to calculate a separate *d*_N_/*d*_S_ for each of the branches of the gene tree. Heterogeneity of *d*_N_/*d*_S_ among branches was tested by comparing the likelihoods of the free-ratios model and model 0, which assumes a homogeneous *d*_N_/*d*_S_ across all sites and branches. This comparison was conducted using a likelihood ratio test [51], assuming that twice the difference between the log-likelihoods of both models 2Δ*ℓ* = 2 × (*ℓ*_FR_ − *ℓ*_M0_), where *ℓ_i_* is the log-likelihood of model *i*, followed a chi-squared distribution with a number of degrees of freedom equivalent to the difference between the number of parameters of both nested models.

To infer the presence of codons under positive selection, we first compared the likelihoods of models M8 and M7. Positive selection was inferred if model M8 (which allows for a class of codons with *d*_N_/*d*_S_ > 1) fitted the data significantly better than mode M7 (which allows *d*_N_/*d*_S_ to vary between 0 and 1). The statistic 2Δ*ℓ* = 2 × (*ℓ*_M8_ − *ℓ*_M7_) was assumed to follow a chi-squared distribution with two degrees of freedom. Next, for each of the branches in the gene tree, a branch-site test of positive selection (Test 2; [42, 43]) was conducted. Positive selection was inferred if model A fitted the data significantly better than null model A1. The statistic 2Δ*ℓ* = 2 × (*ℓ*_MA_ − *ℓ*_MA1_) was assumed to follow a 50%:50% mixture of a point of mass 0 and a chi-squared distribution with one degree of freedom. The Bayes Empirical Bayes approach [42] was used to identify codons under positive selection (posterior probability ≥ 95%).

## ACKNOWLEDGEMENTS

The authors are grateful to Soojin Yi and Julio Rozas for helpful feedback. Computational Resources were provided by Information Technology Operations of the University of Nevada, Reno. This work was supported by a Pilot Grant from the Smooth Muscle Plasticity COBRE of the University of Nevada, Reno, funded by the National Institutes of Health (grant 5P30GM110767-04). MTS was supported by a FPI predoctoral fellowship (BES-2013-062723) and a travel grant (EEBB-I-16-11395) from the Ministry of Economy and Competitiveness of Spain.

## Authors’ contributions

DAP conceived the work and wrote the manuscript. All authors participated in the analysis, and read and approved the final submission.

## SUPPORTING INFORMATION

Table S1: DNMT1 copies in vertebrate genomes.

Table S2: Branch-site tests of positive selection excluding wallaby’s copy 2 and opossum’s copy 1ψ.

